# Ketogenic diet prevents sucrose withdrawal-induced anxiety-like behaviour in male mice

**DOI:** 10.1101/2023.03.10.532006

**Authors:** Mohit Kumar, Roshan Lal, Babita Bhatt, Mahendra Bishnoi

## Abstract

Sugar bingeing has been shown to induce withdrawal effects when sugar is no longer available in the diet. In this study, we investigated the sex differences in sugar-addiction-like behaviour and the effects of the ketogenic diet on these outcomes. We used the two-bottle sucrose choice paradigm as a pre-clinical sucrose overeating and withdrawal model. Female mice consumed more sucrose than males when given free access to water and 10% sucrose for four weeks. One week of sucrose withdrawal after four weeks of consecutive sucrose overeating showed anxiety-like behaviour in male mice. However, the ketogenic diet did not affect sucrose overconsumption in males and females but prevented the sucrose withdrawal-induced anxiety-like behaviour in male mice with a concomitant increase in *cFos* mRNA in the prefrontal cortex. These findings provide evidence of sex differences in sucrose addiction-like behaviour and also indicate that a ketogenic diet may prevent sucrose withdrawal-induced anxiety-like behaviour in males. However, further research is needed to explore the underlying molecular mechanisms driving sex differences in sugar-addiction-like behaviour and the anxiolytic effects of the ketogenic diet.

## Introduction

Sugar bingeing is associated with major health complications including lifestyle diseases (type 2 Diabetes Mellitus and obesity^1^) and co-morbid neurological disorders such as motor learning impairment^2^, anxiety and depression^3,4^. Sugars are rewarding and can recruit neural pathways and mechanisms linked to addiction overriding the eating behaviour from homeostatic eating to hedonic overeating ^5^. Pre-clinical studies have shown that intense sweetness can surpass the cocaine reward^6^. Although the concept of sugar addiction is debatable but has gained much attention recently, whether or not clinical features of addiction can be applied to sugar feeding. Neuroadaptations induced by sugar bingeing lead to severe cravings and withdrawal effects when sugar is no longer available in the diet^7,8^. However, studies exploring the cellular components and underlying molecular mechanisms driving sugar withdrawal symptoms are obscure. Only a few studies have explored the possible underlying mechanisms driving sugar withdrawal associated with negative mood effects. A study by Kim et al. shows that increased levels of Kir2.1 K^+^ channel in dopamine D1 receptors expressing neurons in the nucleus accumbens is responsible for sucrose withdrawal-induced depression and anxiety in mice^7^. Another study by Kawamura et. al. has shown that α7 nicotinic receptor activation suppresses sucrose addiction in mice^8^.

The ketogenic diet (KD), a low carbohydrate, high fat and moderate protein diet, was first introduced in the 1920s by modern physicians as an antiepileptic treatment in children to mimic the fasting metabolism ^9^. KD has gained popularity over the past few years as an effective metabolic therapy for the management of weight gain and hyperglycaemia^10^. In addition to these physiological effects, KD also has the potential to modulate brain functions and behaviour. For example, KD increases locomotor activity^11^ and improves cognitive functions in healthy mice^12^. KD reduces midlife mortality and improves memory in ageing male^13^ and female mice^14^. KD has been suggested as an effective metabolic therapy for the management of anxiety, depression and other mood disorders^15^. A recent study has shown that KD also prevents cognitive decline in mice fed with a high-fat-high-cholesterol diet by suppressing neuroinflammation and improving the expression of brain-derived neurotrophic factor^16^. Altered gene expression^17^, neurotransmitter levels^18^ and metabolic profile of the brain cells^11^ by KD are some suggested molecular mechanisms driving its neuroprotective effects. Recent studies have suggested KD as a potential dietary intervention for the management of ultraprocessed and palatable food addiction^19,20^.

This study was designed with the intent to explore sex differences in sugar-addiction-like behaviour and potential of the KD to prevent these outcomes. Our studies are the first to report sex differences in sucrose addiction-like phenotype and KD as a potential dietary intervention to prevent sucrose withdrawal-induced negative mood effects in males possibly by improving the functionality of the prefrontal cortex.

## Materials and methods

### Animals

Male and female C57BL6/J mice, 6 – 8 weeks of age were used for this study. Mice were procured from the Institute of Microbial Technology (IMTech) Centre for Animal Resources and Experimentation (iCARE), Chandigarh, India, and housed in the Animal Experimentation facility at the National Agri-Food Biotechnology Institute (NABI), Mohali, India. Mice were maintained in a pathogen-free environment at 25 ± 2°C on a 12-h light/dark cycle with food and water *ad libitum* in accordance with the NABI Animal Care and Use Committee. All mice were housed in groups of three to five mice per cage and randomly assigned to treatment conditions after 1 week of acclimatization. Institutional Animal Ethics Committee (IAEC) of NABI approved experimental protocol (NABI/2039/CPCSEA/IAEC/2022/02). Experiments were conducted according to the Committee for Control and Supervision on Experiments on Animals (CPCSEA) guidelines on the use and care of experimental animals. All the experiments performed in the study were blinded at every stage of analysis and the experimenter was unaware of the treatment each animal received.

### Diets and interventions

Normal chow was procured from Hyalasco Biotechnology Pvt Ltd (India). The ketogenic diet (KD; 80% energy from fat) was prepared in-house. The control diet contained (% of total kcal) 18% protein, 65% carbohydrate, and 17% fat. The KD contained 15% protein, 5% carbohydrate, and 80% fat. Milk casein used in the diet was procured from Loba Chemie, India. Animal lard was procured from the local market in Chandigarh, India. Vitamin and mineral mix, DL-methionine and Sodium chloride were procured from Himedia Laboratories Pvt Ltd, India.

### Two-bottle sucrose preference test

Two-bottle sucrose choice paradigm was used in the study to develop sucrose dependence in mice as described previously ^21^. Control mice were given two bottles of water. Sucrose-fed mice were given unlimited access to one bottle each of water (bottle 1) and 10% (w/v) sucrose (bottle 2) for four weeks. Sucrose withdrawal was initiated in half of the animals after four weeks of sucrose overeating by replacing bottle 2 with water and continued for one week. Sucrose consumption, bottle preference and body weight of water and sucrose-treated mice were recorded on 7, 14, 21 and 28 days of the experiment. At the end of the treatment, animals were subjected to behavioural analysis and sacrificed for endpoint biochemical and molecular analysis. The treatment paradigm followed for the study is demonstrated in **Fig. 1**.

**Figure 1.**
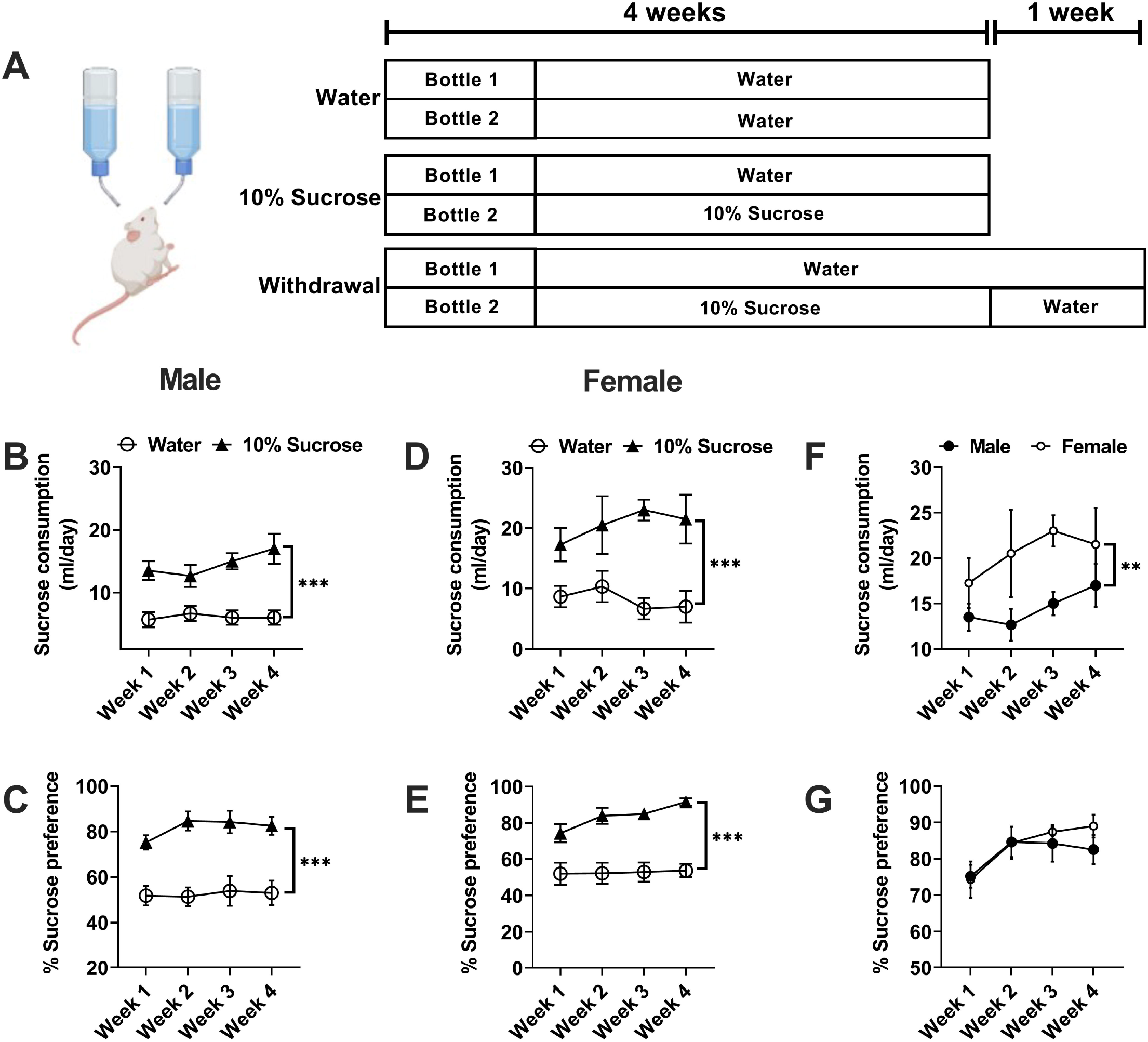
Sex differences in sucrose overeating: Schematic diagram of two-bottle sucrose paradigm followed for the study **(A)**. Time-course analysis of sucrose consumption and sucrose preference in males and females (***p < 0.001, unpaired two-tailed Student’s t-tests; **B-E**). Timecourse analysis of weekly consumption of sucrose in males vs. females (**p < 0.01, unpaired twotailed Student’s t-tests**; F**). Time-course analysis of weekly sucrose preference in males vs. females (**G**). n=10-12 in all treatment groups.

To evaluate the effects of KD on sucrose addiction-like behaviour, two sets of experiments were performed in identical conditions where one set of animals was fed with a control diet and another set was fed with a KD from day 1 to the end of the study. Control or KD-fed mice were given free access to water and 10% sucrose for four weeks. Sucrose consumption and sucrose preference were recorded both in male and female mice. After this, 10% sucrose bottle was replaced with water in the male withdrawal group to initiate spontaneous sucrose withdrawal and was subjected to behavioural analysis and sacrificed for endpoint biochemical and molecular analysis.

### Open field test

The open field test (OFT) was used to assess anxiety-like behaviour in mice. After one week of sucrose withdrawal, all mice were tested for anxiety-like behaviour using the OFT. This test is based on the natural aversion of the animals to explore new areas. Levels of anxiety are directly proportional to the preference of rodents to stay in the central arena. The experimental setup consisted of an arena (50 cm□×□50 cm□×□40.5 cm) placed at a sufficient height to prevent animals to escape. The arena was divided into central and peripheral zones. The mice were placed in the central zone and the activity of each animal was recorded for 15 min using ANY-maze™ tracking software (Stoelting Co. Wooddale, IL, USA). Test chambers were wiped with 70% (v/v) ethanol in between tests to remove any previous scent cues. The ethanol was allowed to dry completely before testing. Parameters such as total distance travelled, freezing time, the number of entries and time spent in the central and peripheral zones were measured.

### Marble-burying test

The marble burying test (MBT) was used to assess anxiety-like behaviour in mice. Mice were placed individually in small cages (29.0 cm x 17.5 cm), in which 20 marbles had been equally distributed on top of mouse bedding (5 cm deep). A lid was placed on top of the cage to prevent the mouse from jumping out of the cage during the test. Mice were left undisturbed for 15 minutes under low light conditions, after which the number of buried marbles (those covered by bedding three-quarters or more) was counted by an observer blinded to experimental conditions.

### Quantitative PCR

Animals were sacrificed by cervical dislocation. The prefrontal cortex was dissected and stored in RNA later (Sigma-Aldrich, St. Louis, MO, USA). Total RNA was isolated using a TRIzol reagent as per the manufacturer’s instructions. cDNA was prepared using RevertAid First Strand cDNA Synthesis Kit (Thermo Fischer Scientific, Waltham, MA, USA) by using 1 μg of total RNA. Primers were designed by using the Primer3Plus primer designing tool. qPCR reactions were assembled using synthesized cDNA, SYBR Green master mix (BioRad Laboratories, Hercules, CA, USA) and 100 nM primers (Eurofins Genomics India Pvt Ltd, Bangalore, India) diluted to 5 nM final concentration. Relative mRNA expression was calculated using the 2^-ΔΔCT^ method. Hypoxanthine phosphoribosyltransferase (HPRT) and glyceraldehyde 3-phosphate dehydrogenase (GAPDH) were used as reference genes for the expression analysis of genes of interest. Sequences of the primers used in the study are provided in **table S1**.

### Statistical analyses

All statistical analyses were performed in GraphPad Prism 8.1.2 (GraphPad Software, La Jolla, CA, USA). Results are presented as mean ± SEM unless otherwise stated. The normality assumption was verified with the Shapiro-Wilk test. Statistical significance was calculated by one-way analysis of variance (ANOVA) followed by the Bonferroni post hoc test for group-wise comparisons unless otherwise stated. Statistical differences between multiple groups were determined using two-way ANOVA with sex and treatment as the primary variables, followed by Bonferroni post hoc test. Outliers were checked for all the treatment groups during the development of graphs.

## Results

### Female mice consume more sucrose than male mice

Male and female mice in the sucrose group consistently consumed a significantly higher amount of sucrose as compared to their water controls during four weeks of sucrose exposure **(Fig. 1)**. Mice given free access to 10% sucrose and water for four weeks consistently showed a higher preference for 10% sucrose (bottle 2) than water (bottle 1), while control animals had no preference. The overall consumption of 10% sucrose was higher in females as compared to males during all four weeks. Male and female mice with unlimited access to 10% sucrose solution did not increase their body weights over the 4 weeks of sucrose exposure **(supplementary Fig. 1)**. Overall this data suggests that females consume more sucrose than their male counterparts when given free access to 10% sucrose and water for four weeks.

### Sucrose withdrawal induces depression and anxiety-like behaviour in male mice

Sugar bingeing has been shown to induce sugar cravings and affective withdrawal symptoms such as anxiety and, negative mood effects when sugar is no longer available in the diet^5,7^. We performed the open field test (OFT) and marble burying test (MBT) after each diet to determine the overall treatment as well as sex-specific effects of sucrose bingeing and sucrose withdrawal on anxiety-like behaviour. In the OFT no overall treatment effects were observed in time spent in the centre zone **(Fig. 2A)**. However, this outcome was sex-specific with only male mice exposed to sucrose withdrawal showing less time spent in the centre zone as compared to their control counterparts. No such differences were detected between any treatment groups in females **(Fig. 2B)**. The observed changes were not due to alterations in the locomotor activity, as no group and sex-specific effects were detected in the total distance travelled in all the treatment groups **(Fig. CD)**.

**Figure 2:**
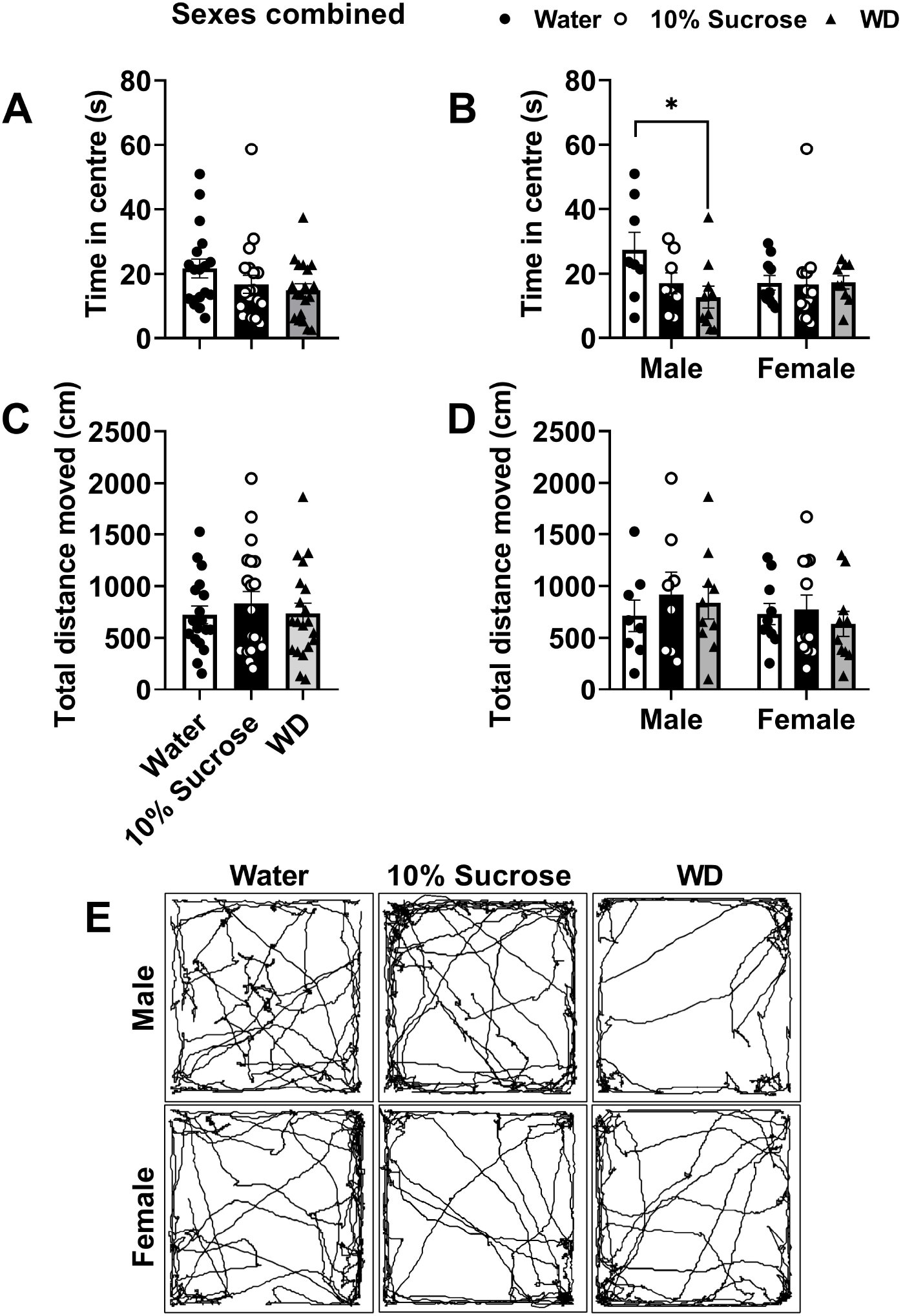
Sucrose withdrawal (WD) induces anxiety-like behaviour in male mice: After 4 weeks of sucrose overeating and I week of subsequent WD, male and female mice were subjected to an open field test to assess anxiety-like behaviour. Bar graph showing time spent **(A-B)** in the centre zone and total distance moved **(C-D)** in the open field test. Track plots of animals in the open field test **(E)**. Data was analysed using two-way ANOVA with Bonferroni post hoc test. (Compared to water- *p<0.05, **p<0.01, ***p<0.001; n=8-20 per treatment group).

In the MBT, post hoc analysis revealed that animals subjected to sucrose withdrawal buried more marbles as compared to their sucrose and water counterparts irrespective of sex **(Fig. 3B)**. Sexspecific analysis showed that only male mice buried more marbles following sucrose withdrawal as compared to their water controls, indicating an anxiogenic-like response **(Fig. 3C)**. In contrast, no differences were observed in the number of marbles buried in water, sucrose, or withdrawal treated females. Overall, our data indicate that sucrose bingeing and sucrose withdrawal induce negative mood effects especially anxiety-like behaviour and these effects are male-specific.

**Figure 3:**
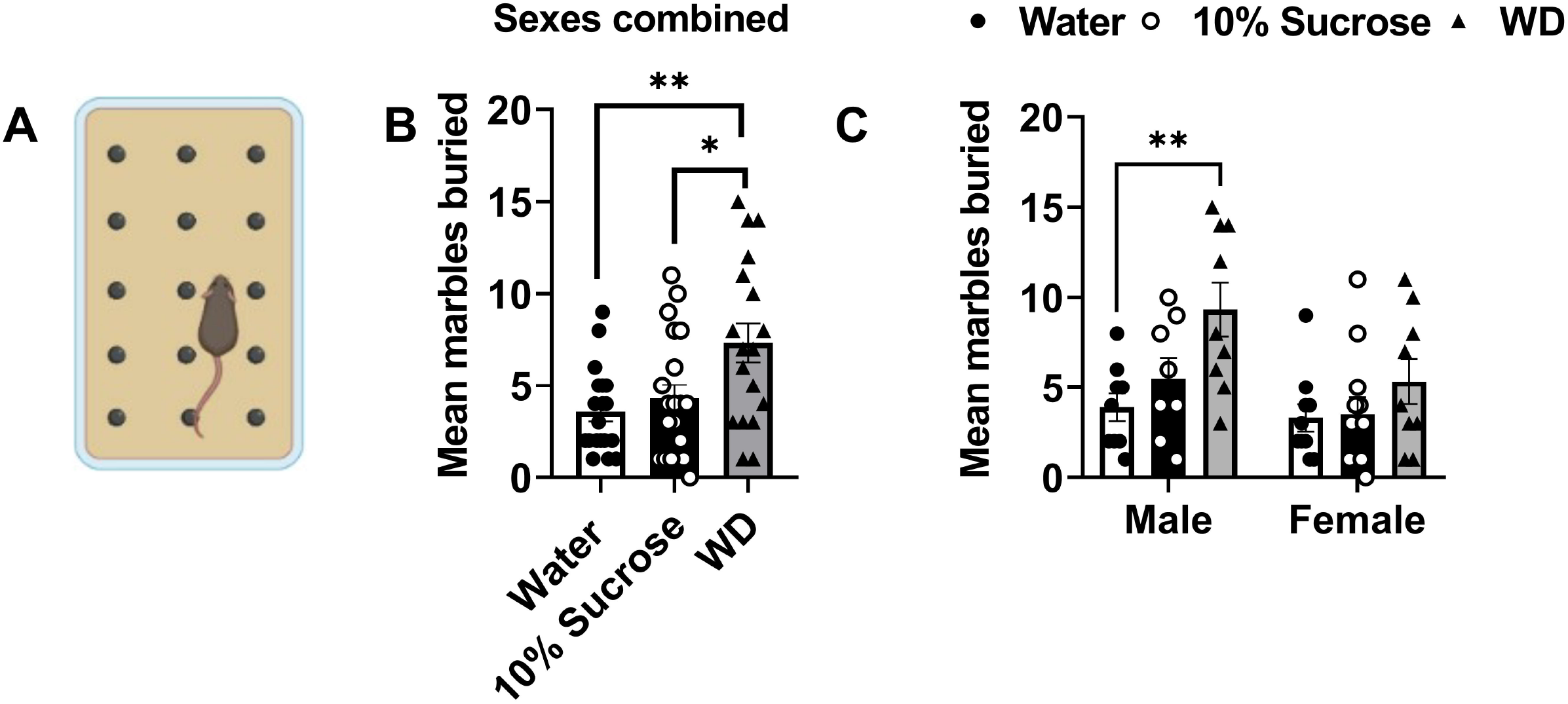
Sucrose withdrawal (WD) induces anxiety-like behaviour in male mice: After 4 weeks of sucrose overeating and I week of subsequent WD, male and female mice was subjected to marble burying tests to assess anxiety-like behaviour. Pictorial representation of the arrangements of marbles in the cage **(A)**. Bar graph showing the number of marbles buried **(B-C)** in the marble burying test. Data was analysed using two-way ANOVA with Bonferroni post hoc test. (Compared to water- *p<0.05, **p<0.01, ***p<0.001; n=8-20 per treatment group).

### The ketogenic diet does not affect sucrose consumption

Recent studies have suggested a low carbohydrate high-fat diet as potential metabolic therapy for the management of binge eating and ultra-processed food addiction^19,20^. Animals were fed with either normal chow or a KD and given free access to two bottles each of water and 10% sucrose for four consecutive weeks. KD did not affect the consumption and preference for sucrose in male and female mice **(Fig. 4)**.

**Figure 4.**
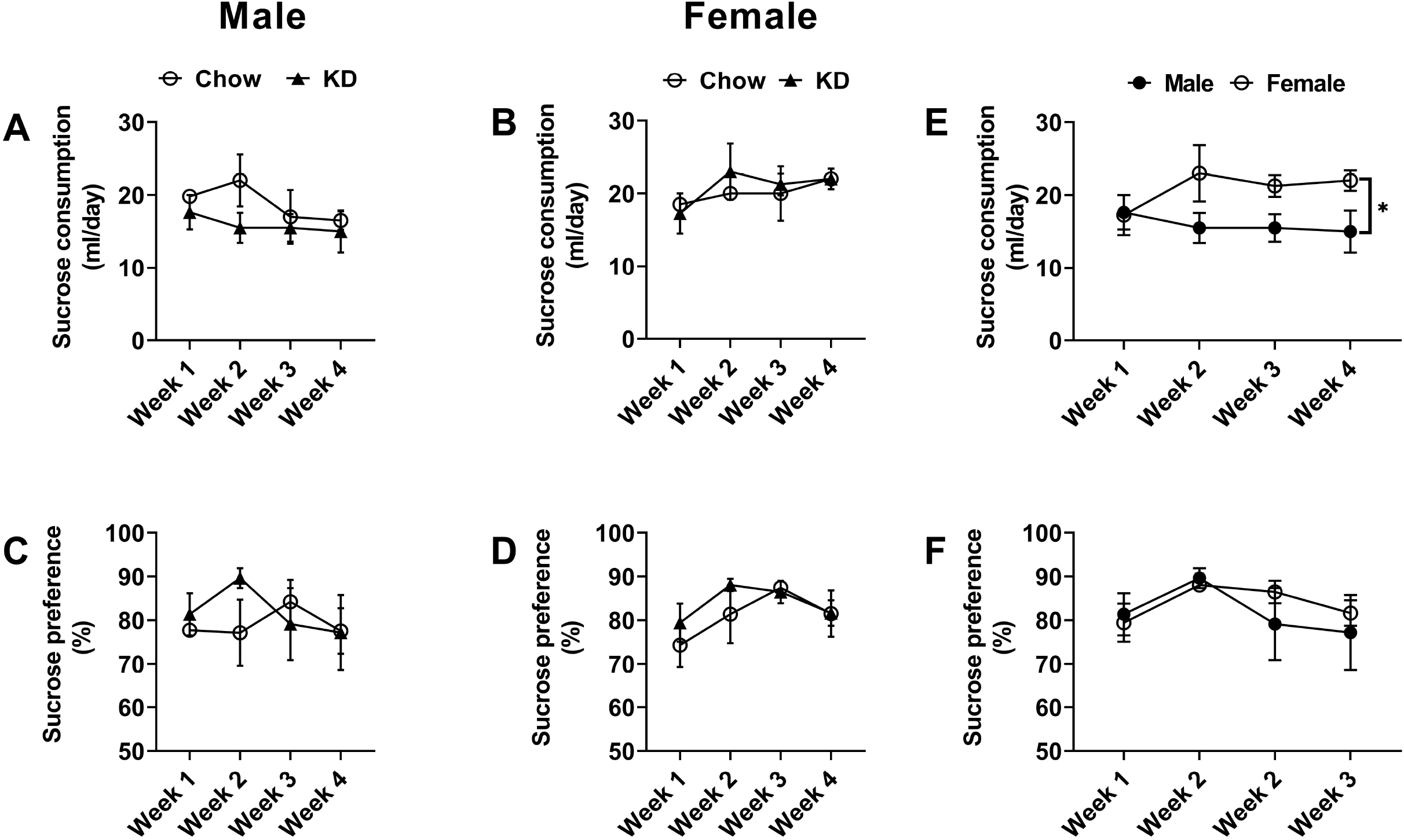
The ketogenic diet (KD) does not affect sucrose consumption in male and female mice. Time-course analysis of sucrose consumption **(A-B)** and sucrose preference **(C-D)** in males and females fed on either a control diet or a KD throughout the study (**B-E**). Time-course analysis of weekly consumption of sucrose in males vs. females fed on a KD throughout the study (*p < 0.05, unpaired two-tailed Student’s t-tests**; F**). n=10-12 in all treatment groups.

### The ketogenic diet prevents sucrose withdrawal-induced anxiety-like behaviour in male mice

Next, we argue if KD can prevent sucrose withdrawal symptoms, especially anxiety-like behaviour when sucrose is no longer available in the diet. To do this, we evaluated anxiety-like behaviour in male mice following sucrose withdrawal fed either with normal chow or a KD. Our OFT analyses showed that control-chow sucrose withdrawal-treated male mice spent lesser time in the centre zone of the arena as compared to their water controls, indicating an anxiogenic-like response; no differences were detected between any groups fed on KD **(Fig. 5)**. The observed changes were not due to alterations in the locomotor activity, as no differences were observed in the total distance travelled between all the treatment groups fed on control chow or a KD (data not shown). In the MBT, sucrose withdrawal-treated male mice on the control chow buried more marbles as compared to their water controls, also indicating an anxiogenic-like response. In contrast, no differences were observed in the number of marbles buried in water and sucrose withdrawal animal groups fed on the KD. Overall, our data indicate that the KD prevents sucrose withdrawal-induced anxiety-like behaviour in male mice.

**Figure 5:**
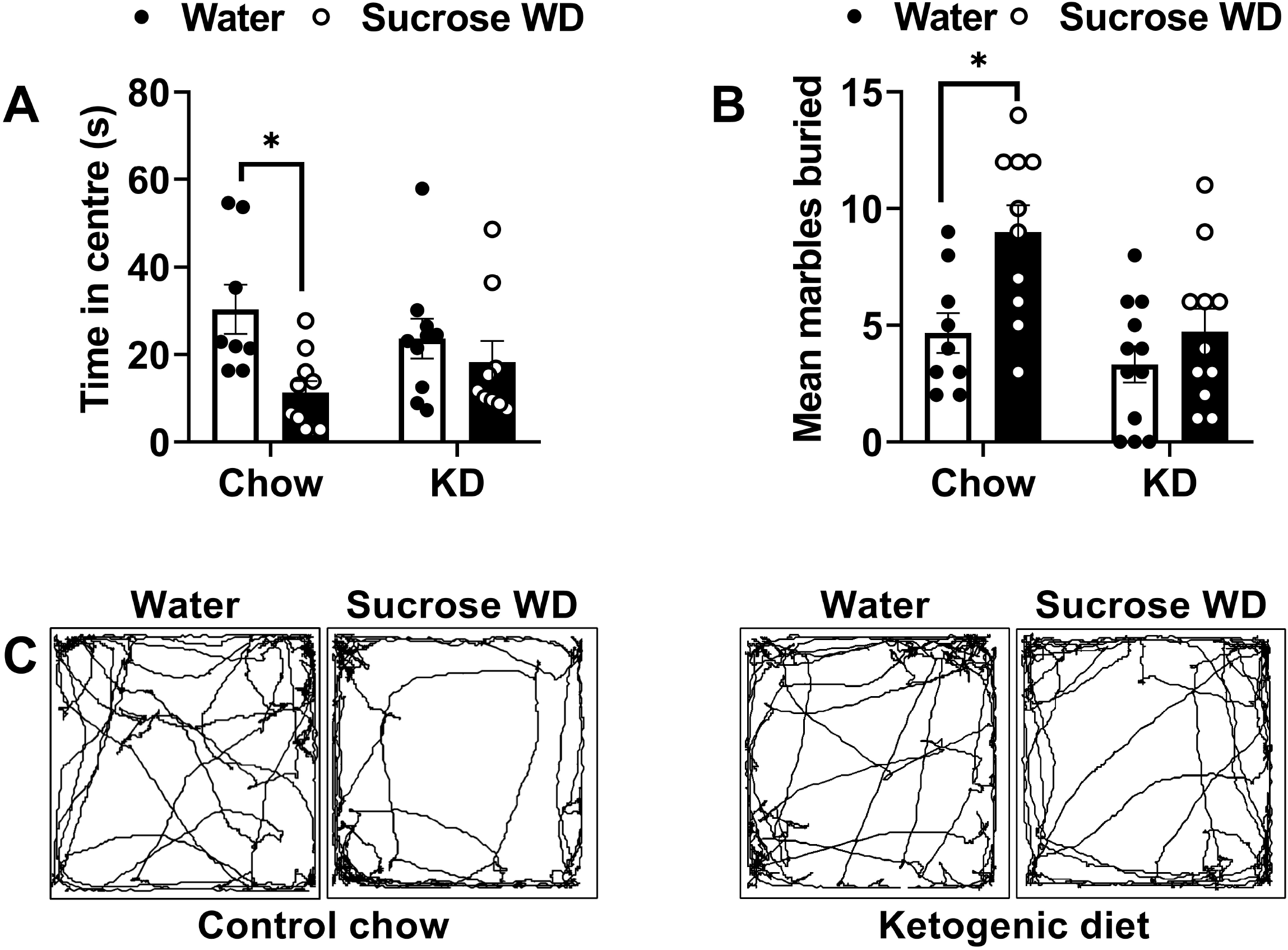
The ketogenic diet (KD) prevents sucrose withdrawal (WD)-induced anxiety-like behaviour in male mice: After 4 weeks of sucrose overeating and I week of subsequent withdrawal, control chow and KD-fed male mice were subjected to the open field test and marble burying test to assess anxiety-like behaviour. Bar graph showing time spent in the centre zone **(A)** in the open field test. Bar graph showing the number of marbles buried **(B)** in the marble burying test. Track plots of animals in the open field test **(E)**. Data was analysed using two-way ANOVA with Bonferroni post hoc test. (Compared to water- *p<0.05; n=8-10 per treatment group).

### Effects of sucrose withdrawal and the ketogenic diet on mRNA expression of *Nox2* and *cFos* in the prefrontal cortex (PFC) of male mice

Our results thus far show that (1) free access to 10% sucrose leads to sucrose overconsumption both in males and females, (2) female mice consume more sucrose than males, (3) supplementation with KD prevents sucrose withdrawal-induced negative mood effects especially anxiety-like behaviour in male mice. Our next steps were to evaluate molecular mechanisms driving the preventive effects of a KD on sucrose withdrawal-induced anxiety-like behaviour in male mice. To do this, we analysed the mRNA expression of NADPH oxidase 2 *(Nox2)* and *cFos* as markers for oxidative stress and neuronal activity respectively in the PFC of male mice **(Fig. 6)**.

**Figure 6:**
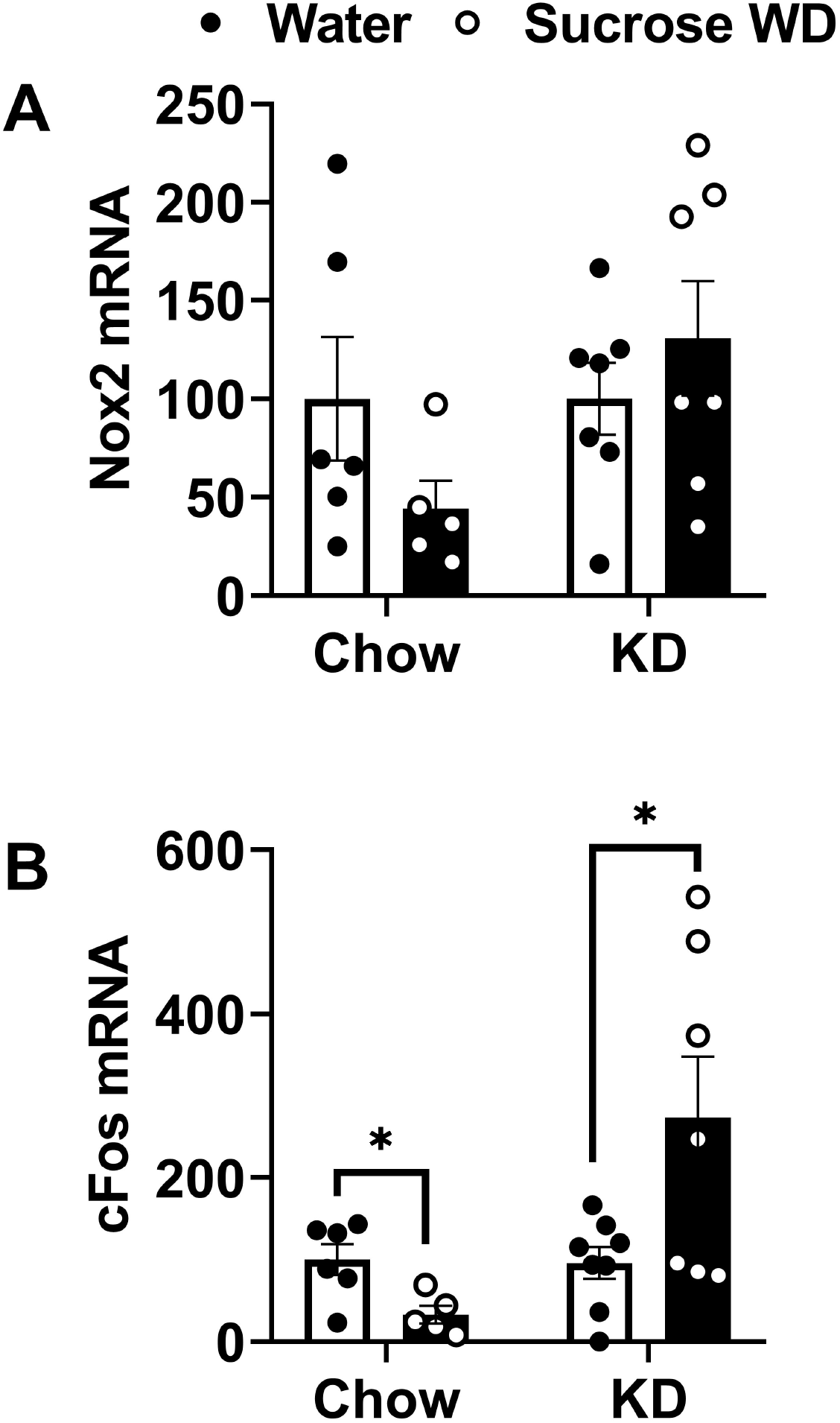
The ketogenic diet (KD) increases cFOS mRNA expression in the prefrontal cortex (PFC) of sucrose withdrawal male mice: Bar graph showing mRNA expression of Nox2 **(A)** and cFos **(B)** in the PFC of male mice. Data was analysed using unpaired two-tailed Student’s t-tests. (Compared to water- *p<0.05; n=8-10 per treatment group).

No change in the mRNA expression of *Nox2* was observed in the PFC of control and KD-fed male mice exposed to sucrose withdrawal as compared to their water controls. mRNA expression of *cFos* was decreased in the PFC of control diet-fed sucrose withdrawal males as compared to their water counterparts. KD-supplemented sucrose withdrawal males showed higher expression of *cFos* mRNA in the PFC as compared to their water controls.

## Discussion

Sugar bingeing in intermittent or regular mode has been shown to induce withdrawal symptoms such as depression and anxiety when sugar is no longer available in the diet^5,7,8^. Dampened dopamine signalling in the nucleus accumbens (primary brain component of the reward pathway) is one of the key factors driving sucrose withdrawal symptoms such as anxiety and depression^7^. These withdrawal effects are responsible for unsuccessful attempts to cut short daily sugar consumption which may lead to obesity and other lifestyle diseases. Here in this study, we demonstrate sex differences in sucrose addiction-like behaviour using a pre-clinical model of sucrose overeating. Our key findings include: 1) Female mice consume more sucrose as compared to male subjects; 2) Sucrose withdrawal induces anxiety-like behaviour in males; 3) The KD prevents sucrose withdrawal-induced anxiety in male mice and these effects are concurrent with increased *cFos* mRNA expression in the prefrontal cortex (PFC). These observations point towards the potential of the KD to prevent sucrose withdrawal-induced negative mood effects, sugar bingeing and susceptibility to develop other lifestyle disorders.

Sex differences are observed in preference, reinforcement and motivation toward sucrose. In operant procedures, female Long-Evans rats consume more sucrose and respond at higher rates for sucrose as compared to males^22^. In another study, female rats consume more sucrose pellets and work harder to obtain a sucrose pellet than males^23^. In a study performed by Datta et. al., male rats show higher motivation toward sucrose than females^24^. In this study, we found that female mice consumed more sucrose as compared to males when given free access to water and 10% sucrose. However, only male mice showed withdrawal symptoms when sucrose was no longer available to the animals after lengthy drinking. Food consumption can be influenced by many factors and food reward is one of the key factors promoting palatable food bingeing. A natural reward is divided into hedonic “liking” and motivational “wanting” behaviour which is regulated by different neural circuits^3,25^. The sex differences in sucrose overconsumption and withdrawal effects we observe in this study are perhaps due to the sex differences in the molecular architecture of the brain components of reward pathways such as the PFC and nucleus accumbens^26^. Sex hormones are also speculated to be involved in the sex-specific response toward sucrose liking and sucrose withdrawal-induced negative mood effects^27^ warranting further research on these lines.

The PFC is one of the primary brain regions engaged in the natural reward process and withdrawal effects^3^. Moreover, neurons in the orbitofrontal cortex of rats are shown to encode compulsion to seek a sucrose reward solution^28^. Acute sucrose withdrawal after prolonged overconsumption may cause transient metabolic and oxidative stress in these brain regions^29,30^. To confirm the hypothesis, we analysed the mRNA expression of *Nox2* in the PFC of male mice. A previous study has shown that Nox2 mediates the high-fat high-sucrose diet-induced nitric oxide dysfunction and inflammation in aortic smooth muscle cells^31^. In contradiction to our hypothesis and the previous study, no significant change in the mRNA expression of *Nox2* was observed in the PFC of the control diet and KD-fed males exposed to sucrose withdrawal. These effects might be cell-specific as a recent study has shown microglia as the primary source of Nox2 in the brain^32^. Therefore, further studies exploring the status of the Nox2 system in the microglia isolated from the brain regions of animals exposed to different diets are urgently warranted.

cFOS is an indirect marker for neuronal activity^33^. Similar to the other drugs of abuse like opioid^34,35^ and alcohol^36^ withdrawal from prolonged sucrose eating might also be associated with dysfunctional PFC as indicated by decreased *cFos* mRNA in male PFC. Activation of endogenous opioid system by sugar^37–40^ might be one of the key factors governing observed sex differences in sugar-induced neuroadaptations and withdrawal symptoms^34^. The expression of *cFos* was restored to normal in the PFC of KD-fed sucrose withdrawal males. KD has been suggested to increase GABA levels^41^ and recent study shows that KD also increases the expression of vesicular GABA transporter (VGAT) in the PFC of mice^42^. Administration of GABA agonist into the PFC has been shown to improve behavioural deficits including anxiety^43^. Hence, GABA-mediated neuronal activation might be one of the mechanisms by which the KD prevents sucrose WD-induced anxiety-like behaviour in male mice. The KD has also been proposed as a potential metabolic therapy for the management of alcohol use disorder^44,45^. It is of great clinical significance to explore the sex- and cell-specific effects of the KD to use this diet for the management of depression, anxiety and substance use disorders.

### In conclusion

findings from this work show sex differences in sucrose addiction-like behaviour and KD as a potential dietary intervention to prevent these outcomes in male mice. These findings will help the susceptible population to adopt a healthy lifestyle by limiting the consumption of sugar and high caloric palatable food further decreasing the chances to develop lifestyle disorders such as obesity in long term. Moreover, these outcomes also emphasize the anxiolytic potential of a KD. Further studies are urgently warranted to explore the possible molecular mechanisms driving high sugar consumption in females, higher susceptibility of sugar withdrawal-induced negative mood effects in males and anxiolytic effects of the KD.

## Author contributions

MK obtained funding for the study. MK designed and performed all the experiments. BB performed real-time RT-PCR studies. RL assisted in mouse behavioural data acquisition. MK wrote the first draft of the manuscript. MB reviewed and provided inputs for the improvement of the manuscript. All authors edited subsequent versions and approved the final version of the manuscript.

## Acknowledgements

Illustrations were created by using BioRender, an online scientific image illustration tool (BioRender.com). We are thankful to Dr Aanchal Aggarwal (NABI) for providing access to ANY-maze™ tracking software (Stoelting Co. Wooddale, IL, USA).

