## Supplementary data for "Ketogenic diet prevents sucrose withdrawal-induced anxiety-like behaviour in male mice"

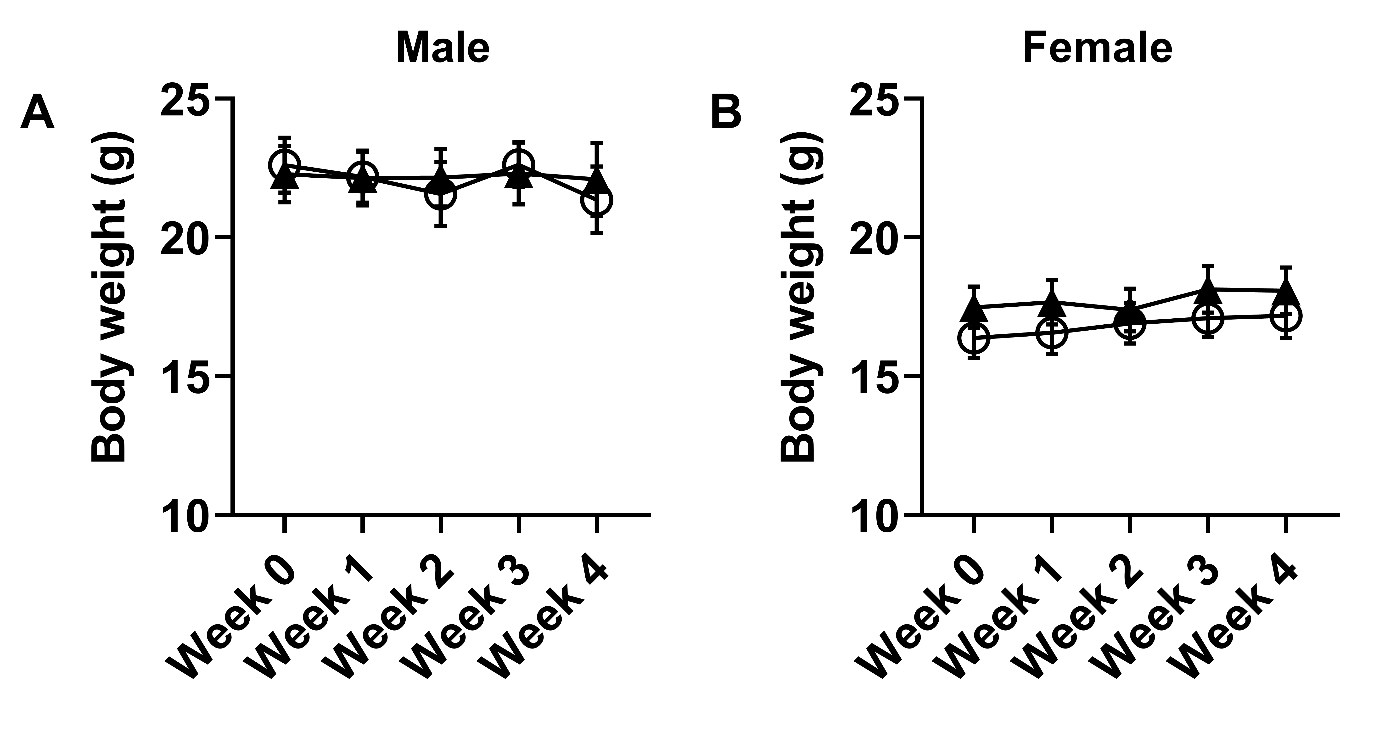

**Supplementary Fig. 1:** Time-course analysis of body weight in males **(A)** and females **(B)**. n=10-12 in all treatment groups.

**Table S1:** Sequence of the primers used for qPCR analysis:

| Gene | Accession Number | Forward Primer | Reverse Primer | Amplicon Size |
| --- | --- | --- | --- | --- |
| *Nox2* | NM_007807 | CCAACTGGGATAACGAGT | GGACATTTGGCAGCATAC | 217 |
| *cFos* | NM_010234 | CTCCCGTGGTCACCTGTA | TTGCCTTCTCTGACTGCTC | 187 |
| *Hprt* | NM_013556 | CAAAGCCTAAGATGAGCG | TTACTAGGCAGATGGCA | 106 |
